# SARS-CoV2 associated secretion of nanoLuciferase reports on virus and Virus-Like Particle production

**DOI:** 10.1101/2022.10.11.511764

**Authors:** Rebekah C. Gullberg, Judith Frydman

## Abstract

SARS-CoV2 is a positive-strand RNA virus in the *Coronaviridae* family that has caused world-wide morbidity and mortality. While much progress has been made we still need expanded rapid anti-virals. The top advanced antiviral candidates all target stages of RNA replication, leaving virus assembly an unexplored avenue of antiviral research. To address this gap, and explore the biochemical and cell biological features of viral assembly, we have employed an improved virus-like particle (VLP) system. We exploited the small nanoLuciferase protein for enhanced signal and surprisingly found that the protein itself appears to be packaged into both SARS CoV2 VLPs and virions and secreted from cells. Interestingly, nLuc is not co-secreted with dengue or Zika infection, suggesting the large virion of Coronavirus can encaspidate and secrete a cellularly expressed reporter protein. Our findings open the way for powerful new approaches to measure viral particle production, egress and viral entry mechanisms.

## Introduction

Severe Acute Respiratory Syndrome Coronavirus 2 (SARS-CoV2) belongs to the family *Coronaviridae*, which are the largest positive-strand RNA viruses that infect humans. SARS-CoV2 is highly transmissible with a broad tissue tropism that is rapidly evolving in human populations. The enormous effort to develop countermeasures against SARS-CoV2 has produced vaccines and antivirals that can reduce viral morbidity, but have limited ability to prevent infection and curtail viral transmission. Research into viral mechanisms necessitates work under cumbersome biosafety level 3 (BSL3) conditions (1,2). The large size of the Coronavirus genome also makes viral reporters difficult to clone and engineer, hindering assays for drug screens.

The importance of developing better sensors and systems to study Coronavirus cannot be understated. In the last 20 years, three separate global coronaviruses outbreaks have spilled over into human populations and have caused extensive morbidity and mortality: SARS-CoV (3,4), MERS-CoV (5) and now SARS-CoV2 (6). These highly transmissible and pathogenic viruses are expected to continue emerging and re-emerging in human populations necessitating expanded rapid tools to improve efforts in diagnostic and therapeutic developments.

Coronaviruses are enveloped viruses with a ∼30kb positive-sense single-stranded RNA genome. Viral particles are ∼100nm in diameter and canonically composed of the spike (S), envelope (E), membrane (M), and nucleocapsid (N) viral proteins (7,8). Despite extensive research, we still have a limited understanding of the mechanisms and pathways of coronavirus viral particle assembly and egress at a cellular level (8,9). Model systems of viral assembly and egress are useful to fill in gaps in our understanding. Virus like particles (VLPs) provide commonly used tools to study viral assembly and egress, permitting studies in BSL2 conditions even for pathogenic viruses. VLPs are composed of viral structural proteins overexpressed in cells that are necessary and sufficient to produce particles that physically resemble the virus of interest, but do not carry the genome, thus are incapable of initiating an infection. VLP systems have applications to understand assembly and egress, as well as entry and neutralization and they are useful as effective vaccine platforms (10,11). Many VLP platforms for SARS-CoV2 have been developed co-expressing the structural proteins in various cell types and ratios as tools for assembly and entry (12–19). A VLP system that harnessed a putative viral RNA packaging signal (PS9) has been recently reported, but is highly inefficient, yeilding 3-5% GFP positive cells after pseudo-infection with concentrated virus (Supplemental Fig. 1). In addition, SARS-CoV2 recombinant systems carrying genetically encoded reporters have been developed, but the large size of the genome makes such efforts cumbersome (20,21). These have been useful to model structural requirements, yet few have demonstrated pseudo-infectivity, indicating that improved platforms are still needed.

Here, we set out to build out a reporter virus-like particle (VLP) system to improve and expand the repertoire of available tools for the more accessible BSL2 research of SARS-CoV2. We exploited the phage-derived MS2 coat protein and MS2-binding loop interaction system to efficiently incorporate reporter molecules GFP and nanoLuciferase into VLPs. Our characterization of this system led us to the surprising finding that the small (19 kDa) nanoLuciferase (herein nLuc) (22) protein itself appears to be packaged into SARS CoV2 VLPs and secreted from cells. Interestingly, nLuc is not secreted upon production of dengue or Zika virus or dengue VLPs, suggesting the large virion of Coronavirus can encaspidate and secrete a cellularly expressed reporter protein. Indeed, we observe that infecting nLuc expressing cells with SARS-CoV2 but not dengue virus leads to secretion of nLuc in a process impaired by inhibitors of viral replication. Our findings open the way to use these two powerful approaches to measure viral particle production, egress and viral entry mechanisms, both in BSL2 (for VLPs) and BSL3 (for WT virus) regimes.

## Results

### Development of SARS-CoV2 virus like particles bearing a reporter mRNA

VLPs are extremely useful as pseudo-infectious reporters for viral entry to screen for inhibitors or to titrate antibody neutralization. Additionally, they mirror authentic virus significantly better than pseudovirus platforms containing only the lentivirus displayed spike protein. Therefore, developing robust reporters of VLP entry is of paramount importance. SARS-CoV2 VLP production by co-expression of the four structural proteins was originally designed for SARS-CoV1 VLP production (23–26) and was used for SARS-CoV2 VLPs (12–19). Based on these, the four canonical structural proteins of SARS-CoV2: spike (S), envelope (E), membrane (M), and nucleocapsid (N) were co-overexpressed in HEK293 cells and the VLPs collected from the supernatant were purified by sedimentation through sucrose cushions followed by analysis on potassium tartrate density gradients (Figure 1A). The migration of structural proteins on density step gradients was examined by immunoblotting and mass spectrometry of gradient fractions. We observed comigration of S, M and N proteins at a region of the gradient where WT virus sediments, consistent with formation of VLPs, E protein was hard to detect, likely because it is very small (11kD), hydrophobic and expected to be packaged at very low levels (14,27,28) (Figure 1B). These analyses indicate VLPs were assembled and secreted from producer cells.

**Figure 1.**
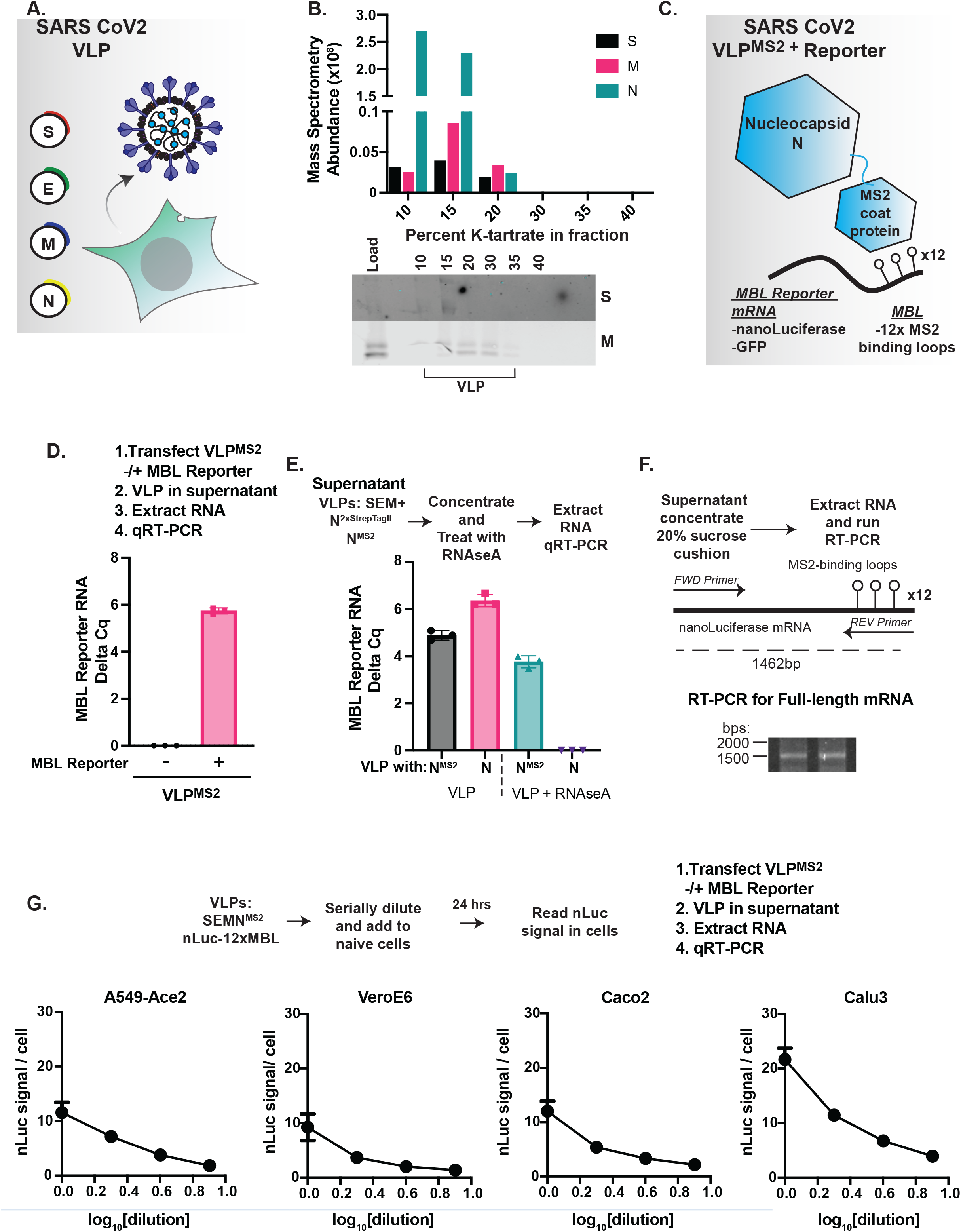
Development of SARS-CoV2 Virus Like Particles with a nanoLuciferase RNA reporter. (A) Schematic of our SARS-CoV2 VLP system. The SARS-CoV2 structural proteins: Spike (S), Envelope (E), Membrane (M) and Nucleocapsid (N) proteins are co-expressed in cells and together form particles that are secreted and resemble virus. (B) <u>VLP proteins are co-secreted from cells.</u> The. supernatnat VLPs were collected, concentrated and run on a density step-gradient. Fractions were collected and analysed by mass spectrometry and western blot. (C) Schematic of trans RNA packaging system. The SARS-CoV2 nucleocapsid (N) protein is genetically fused to the MS2 capsid protein which mediates the specificity of RNA binding. N-MS2 is co-expressed with a plasmid encoding nanoLuciferase (nLuc) and the MS2 binding loops in the 3’ region of its RNA. Together with S, E and M, these are expected to form a virus-like particle encapsulating the reporter RNA, which is expressed upon delivery to new cells. The reporter RNA used here: nLuc or GFP mRNA with 12 copies (12x) of the MS2 binding loops. (D) <u>RNA is secreted with VLPs into the supernatant.</u> VLPs were co-expressed with or without the GFP-12xMBL reporter RNA. qRT-PCR for GFP mRNA show that it is secreted into the supernatant. (E) <u>The N-MS2 protein is responsible for binding the RNA and RNase A protection.</u> VLPs were produced with the N-MS2 construct or N without a tag and the reporter RNA. VLPs were collected, concentrated and incubated with RNase A to degrade any free RNA. RNase A was inactivated with TRIZOL, RNA was extracted and qRT-PCR was run. (F) <u>Full-length mRNA is packaged in VLPs.</u> Schematic for RT-PCR of full-length RNA. VLPs with reporter RNA were concentrated and the RNA was extracted and subjected to RT-PCR with primers to amplify the entire construct. The amplified cDNA was run on a 1% agarose gel. (G). (H) <u>VLPs can enter naïve cells.</u> VLPs with nLuc-12x were serially diluted and added to naïve indicated cells and incubated for 24 hrs and the nLuc signal was measured.

We next tested a recently reported VLP system that incorporates a reporter encoding mRNA using a SARS CoV2 viral packaging signal system (12). Upon addition of these VLPs to Ace2-expressing cells, we observed the encoded Firefly Luciferase (FfLuc) yielding activity in the recipient cells, but the very low signal level limits the dynamic range of measurements (Supplemental Figure 1). Therefore, we set out to build a stronger reporter system for SARS-CoV2 VLPs. We reasoned that the bacteriophage capsid MS2-RNA binding interaction, which has well validated specificity and higher affinity, could enhance the ability of VLPs to incorporate more copies of reporter mRNA into the particle, yielding an improved signal to background ratio upon delivery to recipient cells. As reporters, we chose GFP and nanoLuciferase (nLuc) as reporter molecules. Similar to FfLuc, nLuc has a facile and sensitive luminescence assay (29). Our VLP design consisted of the four SARS-CoV2 structural proteins whereby where the N protein carried a C-terminal fusion to the MS2 bacteriophage capsid protein (N^MS2^ (30)) as well as a reporter gene containing 12 copies of the MBVSV6 loop corresponding to an improved MS2 bacteriophage packaging signal (30,31) (Figure 1C). We expected that the MS2 capsid domain in N^MS2^ will recruit the mRNA carrying the MS2-binding loops (herein MBL), to yield a VLP carrying the encoded reporter in the viral particle (herein VLP^MS2^).

### The reporter RNA is packaged and secreted with SARS-CoV2 VLPs in an MS2-dependent manner

To test if the reporter RNA can be found in secreted VLP^MS2^ we used qRT-PCR to measure GFP RNA levels the supernatant. When transfected with the VLP^MS2^, we detected the GFP mRNA in the supernatant for GFP (Figure 1D and E). To ensure that the secretion of the RNA is dependent on recruitment by N^MS2^, we co-transfected the nLuc reporter with the structural proteins that generate either VLP^MS2^ or VLP without MS2 protein domain. We collected the secreted VLPs and treated them with RNase A, to digest any mRNA that is not encapsulate. Following RNase A inactivation, we extracted the RNase-resistant RNA using Trizol and examined the levels of reporter RNA by qRT-PCR. Only the presence of N-MS2 protein yielded secreted mRNAs that were protected from degradation, consistent with specific recruitment into VLP^MS2^ (Figure 1E).

We next examined whether the VLP^MS2^ particles package and secrete the full-length mRNA for the reporter. Thus, we designed primers to amplify the entire expected RNA construct: ∼1400bp of the nLuc and 12xMBL loops (Figure 1F). Using these primers, we amplified RNA from purified VLPs with a longer extension time to make cDNA, followed by agarose gel analysis. This revealed VLPs package full-length RNA as expected for mRNA competent as a reporter (Figure 1F). This does not rule out the possibility of VLPs also incorporating smaller RNA molecules.

Finally, we tested the ability of our reporter VLP^MS2^ bearing nLuc to pseudo-infect new cells. We serially diluted these VLPs and added them to a panel of ACE2 expressing cells, allowed them to attach for 1hr, washed the inoculum and added fresh media. After a 24 hr incubation, cells were lysed and we quantified nLuc activity (Figure 1G). For all the cell types, we observed found robust dose-dependent nLuc signal, indicating the utility of the VLP^MS2^ reporter system.

### Packaging and secretion of nanoLuciferase protein by SARS-CoV2 VLPs

Motivated by our findings that the reporter RNA is packaged and delivered to naïve cells, we set out to determine the efficiency of VLP based delivery of RNA through direct ‘infection’ compared to direct transfection. The supernatant of cells producing VLPs carrying the nLuc reporter RNA was collected and filtered. A fraction (0.1 ml) was directly added to naïve HEK293T-Ace2/TMPRSS2 cells, while 1 ml of the same supernatant was used to extract mRNA which was transfected into naïve cells (Figure 2A). After 24 hrs we measured the nLuc signal in the recipient cells. Both approaches led to nLuc expression in the naïve cells (Figure 2A). However, the nLuc signal from the direct ‘infection’ was significantly higher than that obtained by transfection of RNA, even when extracted from a larger fraction of supernatant.

**Figure 2.**
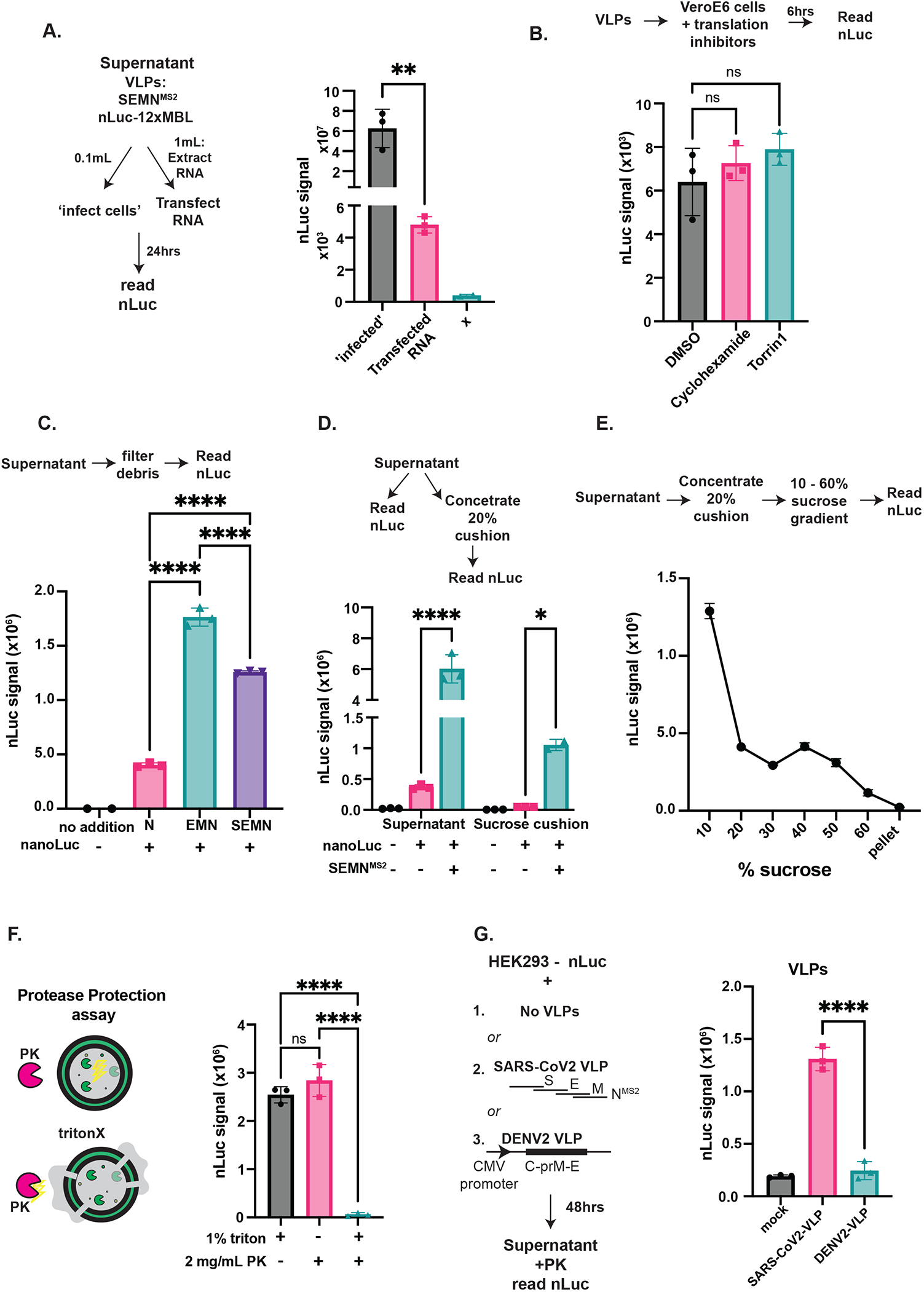
nanoLuciferase protein is packaged and secreted from VLP expressing cells. (A) Schematic of ‘infection’ and transfection approaches to deliver reporter VLPs to new cells. VLPs with the nLuc-12x construct were produced and RNA was extracted from the supernatant. The supernatant was added directly to naïve cells, while other naïve cells were transfected with the RNA. nLuc signal was read after 24 hrs of incubation. (B) <u>The nLuc signal in naïve cells is not dependent on cellular translation</u>. Reporter VLPs were added to naïve cells and treated with cycloheximide or torrin1 for 6hrs to prevent any translation of the reporter message. We see no difference in nLuc signal indicating that the signal does not arise from nascent translation of the reporter RNA. (C-D) <u>nLuc protein is co-secreted with VLPs</u>. The nLuc-12x reporter along with the indicated VLP proteins were transfected into cells and incubated for 48hrs. The supernatant was removed, filtered and the nLuc signal was measured directly in the supernatant. (D) Similar supernatant was processed for nLuc measurement and also concentrated through a 20% sucrose cushion and nLuc signal was measured in the pellet. (E) <u>nLuc is packaged and secreted with VLPs</u>. VLP supernatant was again concentrated through a 20% sucrose cushion and then separated on a sucrose density gradient and nLuc was measured throughout the gradient indicating nLuc associating with high molecular weight species. (F) <u>VLPs package nLuc.</u> Schematic of protease protection assay. VLPs were produced with nLuc and concentrated through a 20% sucrose cushion. The purified VLP sample was then left untreated or treated with 1% triton followed by protease K (PK). Samples treated with only PK were protected, while the nLuc signal was removed by PK in the triton disrupted samples, indicating that a majority of the nLuc is packaged in a vesicle (G) <u>nLuc is not packaged into DENV2 VLPs</u>. HEK293 cells were transfected with nLuc and GFP or SARS-CoV2 or DENV2 VLP proteins. The supernatant was collected after 48hrs, filtered, treated with PK and nLuc signal measured. One-way ANOVA with multiple comparisons test was performed: ns=not significant, *=p<0.05, **=p<0.01, ***p<0.001, ****p<0.0001; n=3.

Our experiment suggested VLP based delivery of nLuc was more efficient than traditional lipid-based transfection methods. To ensure the nLuc activity in VLP “infected” cells arouse from mRNA translation, we examined the effect of two different protein synthesis inhibitors, cycloheximide and torin, added after addition of VLPs to naïve cells. Surprisingly, translation inhibitor did not diminish the nLuc signal from VLP addition (Figure 2B). Therefore, we considered whether the VLPs contain nLuc protein in addition to the reporter mRNA, and whether this contributes substantially to the signal VLP “infection”. Indeed, the filtered supernatant from cells expressing nLuc and VLPs (Figure 2C) as well as VLPs purified through a sucrose cushion (Figure 2D) both contained nLuc activity that could be measured directly. This indicates that nLuc is co-secreted from cells in a VLP-dependent manner.

We next determined if nLuc is actually packaged inside the VLPs or if it is co-secreted as a free protein. First, we subjected the sucrose-cushion purified VLP-nLuc to fractionation on a sucrose density gradient and we measured nLuc activity in each fraction. Indeed, a fraction of nLuc was found in a large structure, migrating in similar density fractions as the VLPs (Figure 2E). Secondly, we used a protease protection assay to determine if nLuc is contained within a lipid enveloped structure (Figure 2F). Indeed, most nLuc signal was protected from digestion with Proteinase K (PK). However, nLuc activity was lost when the PK treatment was carried out after treatment with 1% triton to disrupt the envelope, consistent with nLuc being contained within a VLP (Figure 2F).

To determine if other VLP systems also promote nLuc secretion, we compared the SARS-CoV2 VLPs system to a DENV2 VLPs system where the capsid, prM and envelope proteins are co-expressed in a single plasmid (32) (Figure 2G). Cells expressing nLuc were transfected with either the SARS-CoV2 VLP or the DENV2 VLP plasmids, and the VLPs collected and filtered and treated with PK to remove any soluble nLuc protein. We then measured the nLuc activity in the treated VLP. We found robust secretion of nLuc with SARS-CoV2 VLPs but not with DENV2 VLPs (Figure 2G) suggesting a SARS-CoV2 specific effect.

### SARS-CoV2 virus but not dengue or Zika viruses packages nanoLuciferase reporter

We next asked whether the SARS-CoV2 virion can package nLuc similarly to the VLPs., We thus infected A549-Ace2 cells (Figure3A) or Hek293T-A2T2 cells (Figure 3B) that were expressing nLuc with SARS-CoV2 using two multiplicity of infections (MOIs). After 24 hrs, we collected both the cells and their supernatant and measured nLuc activity in the presence and absence of virus infection. nLuc was expressed in the cytosol of all cells but the nLuc signal in the supernatant was increased upon viral infection. As observed for the VLPs, a PK protease protection assay followed by an nLuc activity assay indicated the nLuc secreted from cells upon SARS-CoV2 infection was protease resistant, but was rendered PK sensitive upon treatment with 1% triton, suggesting that nLuc is packaged in a protease protected, triton sensitive vesicle during viral infection (Figure 3C).

**Figure 3.**
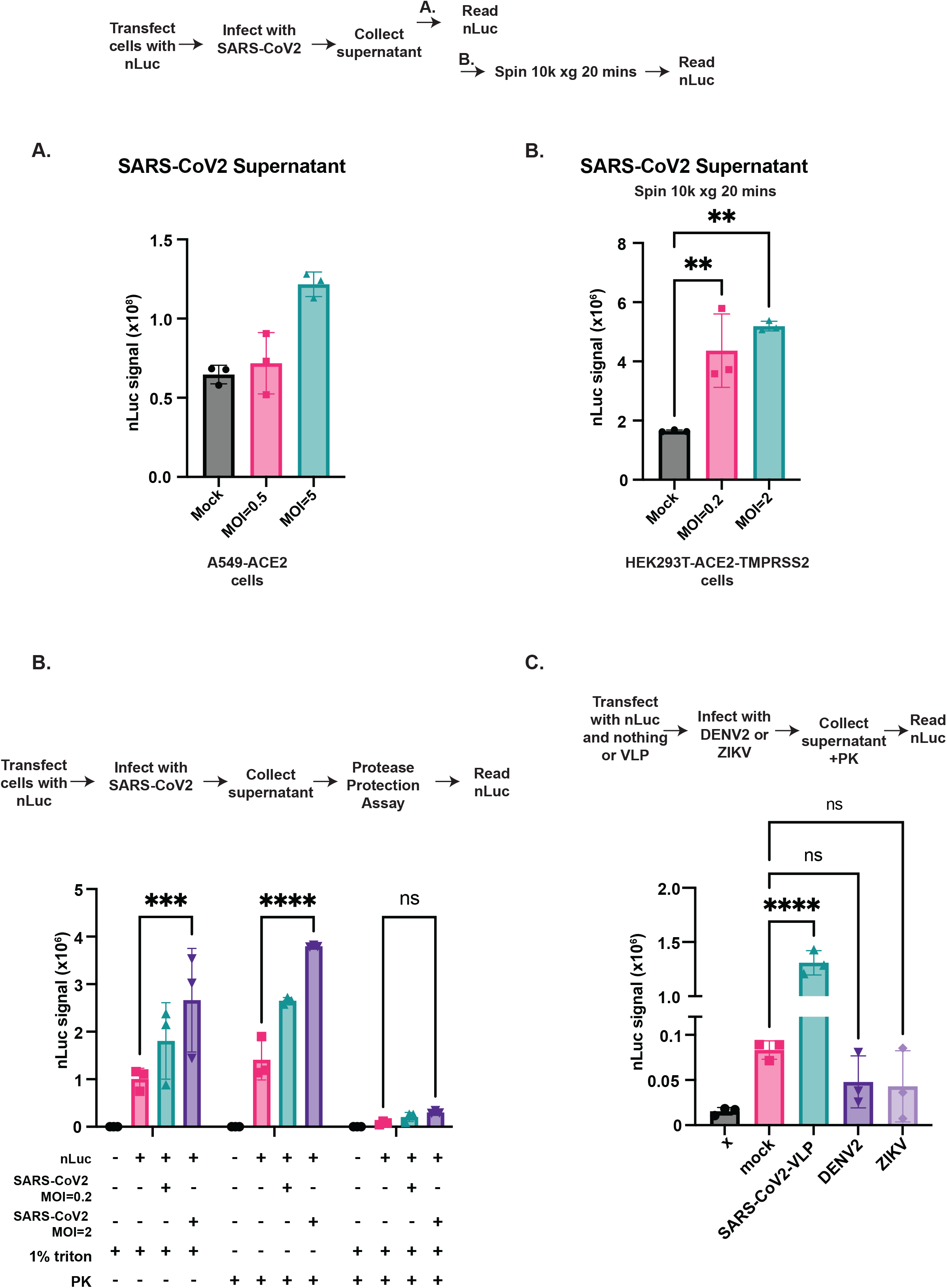
SARS-CoV2 induces packaging and secretion of nanoLuciferase. (A) <u>nLuc in secreted in virus-infected cells</u>. A549-ACE2 cells were transfected with nLuc and infected with SARS-CoV2 for 24 hrs and the cells and supernatant were directly read for nLuc signal and the secreted signal was normalized to cell expression. (B) Hek293T-ACE2/TMPRSS2 were transfected with nLuc and infected with SARS-CoV2 for 48 hrs. The supernatant was collected and debris were spun down at 10kxg for 20 mins and then nLuc was measured. (C) <u>SARS-CoV2 induces packaging and secretion of nLuc</u>. nLuc transfected cells were infected with SARS-COV2 and the supernatant was collected and clarified and treated with 1% triton and PK. We see that with a higher MOI more nLuc signal remains protected from PK treatment indicating virus-induced packaging in secreted vesicles. (D) <u>nLuc is not packaged into DENV2 or ZIKV virus particles</u>. HEK293 cells were transfected with nLuc and infected with DENV2 or ZIKV. The supernatant was collected at 48hrs, filtered treated with PK and nLuc signal measured. One-way ANOVA with multiple comparisons test was performed: ns=not significant, *=p<0.05, **=p<0.01, ***p<0.001, ****p<0.0001; n=3.

To assess the generality of our findings for other enveloped viruses, we infected nLuc expressing cells with two flaviviruses: DENV2 strain 16681 (33) or ZIKV (PRVABC59) (34) at MOI=0.5. We utilized the SARS-CoV2 VLP as a positive control. We collected the supernatant at 24 or 48 hrs and examined the levels of PK-resistant nLuc activity in the supernatant. While the SARS-CoV2 VLPs, which served as a positive control, did contain nLuc activity, there was no nLuc signal with either DENV nor ZIKV at either timepoint. Thus, nLuc secretion is not a common feature of all viral infections, but it does occur in SARS-CoV2 (Figure 3D and Supplemental Figure 2).

### Secreted nanoLuciferase reflects SARS-CoV2 replication levels and can be used as a screening tool

To establish a link between SARS-CoV2 virus production from cells and nLuc secretion we examined the effect of siRNA knockdown of the viral RNA-dependent RNA polymerase (RdRP), an intervention that reduces viral replication. We then compared nLuc particle secretion to secreted viral RNA copies and infectious titer. Cells expressing nLuc were treated with either non-targeting siRNA or an siRNA targeting the SARS-CoV2 RdRP (35) and the infected at MOI=0.5. After 48 hrs we collected and processed the supernatant and measured RNA copies via qRT-PCR, infectious virus titers via plaque assay and nLuc signal (Figure 4A). We found a significant decrease in the number of secreted RNA copies of RNA as well as a decrease in infectious virus titers with RdRp knockdown as expected (Figure 4B). We also observed a significant decrease in secreted nLuc upon siRNA RdRP treatments (Figure 4C). The decrease in nLuc signal mirrors the decrease in viral RNA copies with siRNA treatment as well as infectious virus titers (Figure 4D). Therefore, packaged and secreted nLuc can be used as a rapid, useful and relevant tool to measure virus associated secreted particles.

**Figure 4.**
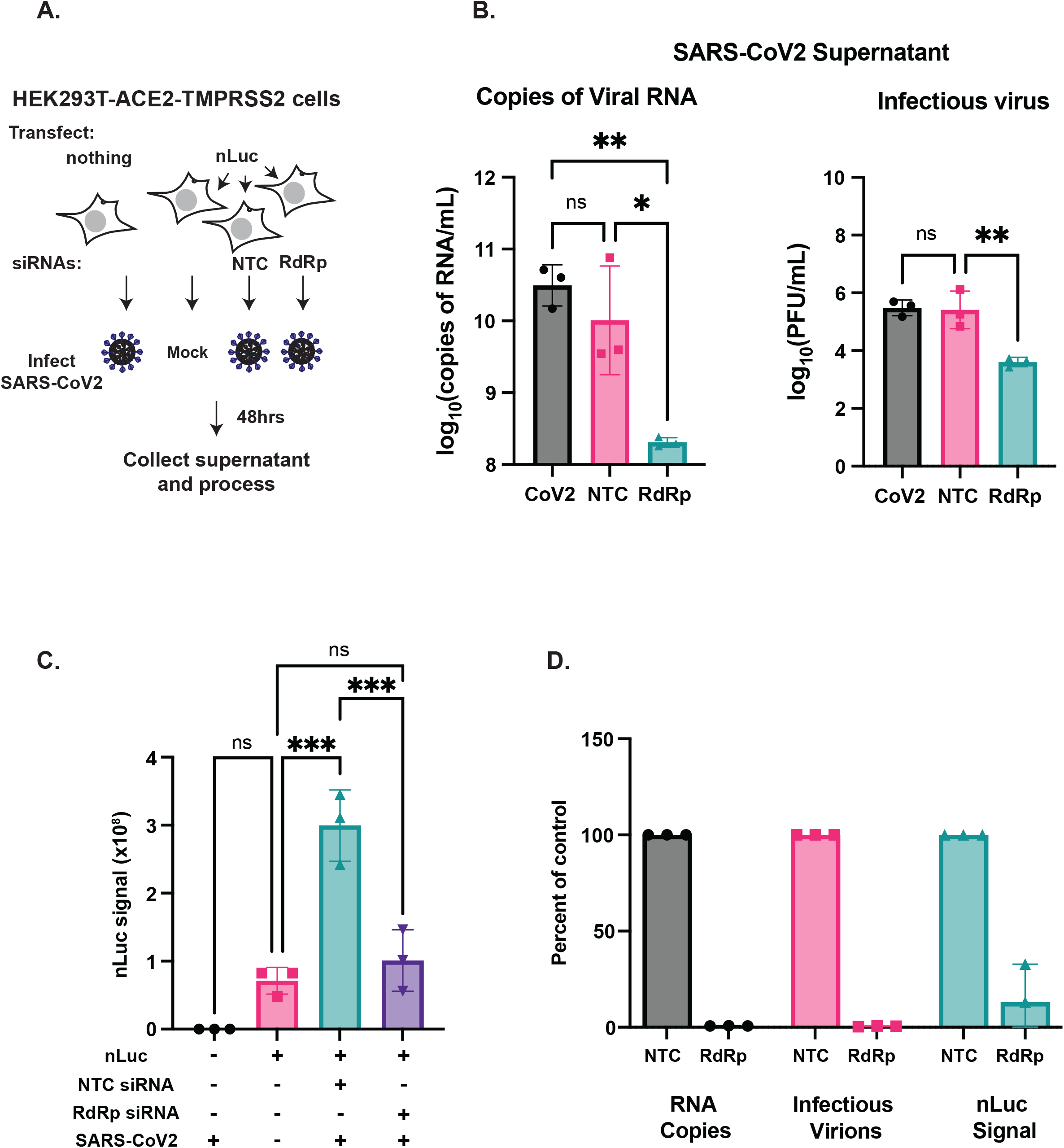
SARS-CoV2 associated secreted nanoLuciferase mirrors viral particle secretion. (A) Schematic of cells transfected with nLuc and siRNAs with viral infection. (B) <u>Knockdown of SARS-CoV2 RdRp reduces infectious virus and particle formation</u>. HEK293T-ACE2/TMPRSS2 cells were treated with siRNAs targeting the RdRp or a non-targeting control. The cells were subsequently infected with SARS-CoV2, MOI=0.5 for 48 hrs. The supernatant was collected and processed for viral genome equivalents, and infectious virus via plaque assay. (C) nLuc was measured in the same supernatant (D) <u>nLuc secretion correlates with secretion of total and infectious viral particles.</u> The percent of control was measured for RNA copies, PFU and nLuc signal with RdRp knockdown compared to non-targeting control. A One-way ANOVA with multiple comparisons test was performed: ns=not significant, *=p<0.05, **=p<0.01, ***p<0.001, ****p<0.0001; n=3.

## Discussion

Here, we have demonstrated a novel method to rapidly measure secreted SARS-CoV2 viral particles that mirrors secreted RNA and infectious virus. While we set out to produce a novel reporter RNA construct useful for VLP entry assays, we found the true indicator masked by background signal rendering it non-specific as a screening tool for spike-ACE2 dependent entry. However, by exploring of the molecular nature of this surprising background signal, we discovered a novel phenomenon of SARS-CoV2 viral particle assembly that can be exploited it as a tool to measure viral particle secretion.

While our reporter RNA particles did package full length reporter mRNA that can be translated in recipient cells, its pseudo-infectivity based activity is close to background levels. Only one other SARS-COV2 VLP system has demonstrated spike-ACE2 dependent pseudo-infectivity, but also in that system only 3-5% of cells are ‘infected’ with VLPs (12). Therefore, proper SARS-CoV2 VLP assembly is an inefficient process in the absence of viral infection.

Surprisingly, nLuc protein itself is secreted in a protease protected, triton sensitive particle during both VLP expression or SARS-CoV2 viral infection. Interestingly, we also observed that the uninfected cells (mock) have packaged and secreted nLuc. This may be a phenomenon of overexpression and secretion of nLuc in some type of extracellular vesicle. It could be the case that viral infection enhances this process and increases the secretion of extracellular vesicles with nLuc or it could be that the nLuc is also packaged inside viral particles. Further biochemical analysis of the secreted vesicles is in order to determine if they are authentic viral particles or co-secreted vesicles.

The packaging and secretion of nLuc that we observe here, although specific to SARS-CoV2, occurs through an undefined mechanism. We hypothesize that the mechanism is mediated through the molecular sizes of nLuc and SARS-CoV2. DENV2 and ZIKV are flaviviruses that make small viral particles, approximately 50nM in diameter, which bud into the lumen of ER membranes (36). SARS-CoV2, however generates particles that are approximately 100-150nM in diameter (7,8), nearly three times larger than flaviviruses. The large size of coronaviruses is likely to accommodate their large genomes; however, it also suggests the possibility for flexibility in the cargo-carrying capacity of coronaviruses and the ability to incorporate extra small proteins. Thus, since cytoplasmic nLuc is abundant when overexpressed in cells, it is readily packaged into viral particles as they are incorporating their cytoplasmic nucleocapsid into a budding particle. This works lays the foundation for future work to characterize what else may be packaged into viral particles as well as the specificity of the budding process.

Rapid tools to measure secreted viral particles are needed to address our current and future pandemics. Here we demonstrate a rapid assay that is specific to SARS-CoV2 VLP and virions to measure the secreted total particles that tracks with secreted RNA copies and viral infectivity. It is readily adaptable to future variants of SARS-COV2 as well as other novel coronaviruses.

## Methods

### Plasmid production

The codon optimized plasmid library of the SARS-CoV2 viral structural proteins was a gift from Nevan Krogan (37). The N proteins was cloned into new vectors to produce the N-MS2 for use in this study. NanoLuciferase and EGFP were cloned into constructs with 12 copies of the V6-MBL (31). The Dengue virus 2 VLP plasmid was a kind gift from Dr. Ted Pierson (32).

### Expression and purification of mammalian VLPs

#### Cells

VLPs were produced in adherent HEK-293 [HEK-293](ATCC-CRL-1573) or the suspension Expi293 cells Expi293 (Gibco). HEK-293 cells were maintained in 10% fetal bovine serum (Atlanta Biologicals) in Dulbecco’s modified Eagle’s medium (DMEM; Gibco) with 1% penicillin–streptomycin (Gibco) and 1% non-essential amino acids (neAA; Gibco) at 37°C in a humidified incubator with 5% CO_2_. Expi293 cells were maintained in Expi293 Expression Medium (Gibco) at 37°C in a humidified incubator with 8% CO_2_ shaking at 130 revolutions per minute.

#### Transfection

HEK-293 cells were transfected with the indicated plasmids mixed at a 1:2 (µg DNA : µL transfection reagent) ratio with Lipofectamine2000 (Invitrogen) in OpitMEM (Gibco) following the manufactures protocol. The Expi293 cells were transfected with the indicated plasmids at a 1:2 (µg DNA : µL transfection reagent) ratio with the ExpiFectamine 293 reagent following the manufacturer’s instructions.

#### VLP collection

The VLP producing cell supernatants and cell pellets were collected after 48hrs and processed. The supernatants were clarified at 1000xg for 10 mins at room temperature, then filtered through a 0.45uM PVDF membrane (Milipore Sigma) to remove cell debris.

#### VLP concentration and purification with sucrose cushion ultracentrifugation

To concentrate and purify proteins in a higher density complex such as a VLP, the samples were pelleted through a 20% sucrose in 120 mM HEPES cushion at 100,000xg for 2hrs at 4 °C with 45 Ti or 70 Ti fixed-angle rotor (Beckman-Coulter) depending on the volume of the sample. After centrifugation, the supernatants were removed, the ultracentrifuge tubes were inverted for 5 minutes on a paper towel to remove remaining supernatant and the pellets were gently resuspended in TNE buffer overnight at 4°C and equal volume was processed.

#### Gradient purification of VLPs

To biochemically characterize the density of the VLPs and further purify them we ran Potassium Tartrate density gradients. Briefly, after concentration and an initial purification through a 20% sucrose cushion the resuspended pellet of VLPs was loaded on top of a hand-poured step-gradient consisting of top to bottom: 5 - 40% K-tartrate with 3.75-30% glycerol in TNE buffer. The gradients were centrifuged at 200,000xg for 3 hs at 4°C in a SW-41Ti rotor. The gradients were carefully removed and fractions collected from the top by slowly pipetting 1 mL fractions. Each fraction was then buffer exchanged into TNE by centrifugation through a 100kD 0.5mL amicon filter.

### Western blotting

To quantify structural proteins in the gradient, 10 μl of VLPs were denatured with 2x leamlli buffer (2%SDS, 10%glycerol, 0.002% bromophenol blue and 0.06125MTris-HCl) with 5% β-mercaptoethanol at 50°C for 10mins and loaded onto a 4-20% Tris-glycine SDS-PAGE gel (Invitrogen). The proteins were transferred to a nitrocellulose membrane with an iBlot (ThermoFischer), which were blocked with 3% BSA in TBS containing 0.2% tween-20 (TBS-T) and probed with rαstrep (Abcam, polyclonal, ab76949), overnight at 4°C with gentle rocking. The membranes were then washed 3 times in TBS-T and incubated with IRE-dye-800 - conjugated anti-rabbit antibody at 1:10,000 dilution for 1 h, washed again 3 times in TBS-T followed by 3 washes in DI water and imaged on the LiCor scanner.

### RNA extraction and qRT-PCR

RNA was extracted from cells or supernatant using Trizol or Trizol LS (ThermoFisher) respectively. A one-step qRT-PCR kit with SYBR green (Agilent) was used with the primer sequences below. Reactions were set up according to the manufacturer’s protocol and run on a BioRad CFX96 Touch Real-Time PCR Detection System. The cycling parameters were: 20 mins at 50°C for reverse transcription, then 5 mins at 95°C followed by 45 two-step cycles of 95°C for 5 seconds and 60°C for 60 seconds, followed by a melt curve starting at 65°C and ending at 97°C. The annealing/extension step was extended to 5 mins at 60°C for the full-length RT-PCR. Primers were designed using Benchling primer design software for nsp15, EGFP, full-length nLuc-12xMBL, and NCBI for GAPDH. A standard curve of *in vitro* transcribed RNA from a SARS-CoV2 cDNA subclone (nucleotides: 19785-20364) was generated and used to quantify the viral genome copies in the supernatant. Delta Cq values were calculated by subtracting the measured Cq value from a blank control or a cut off of 30 cycles.

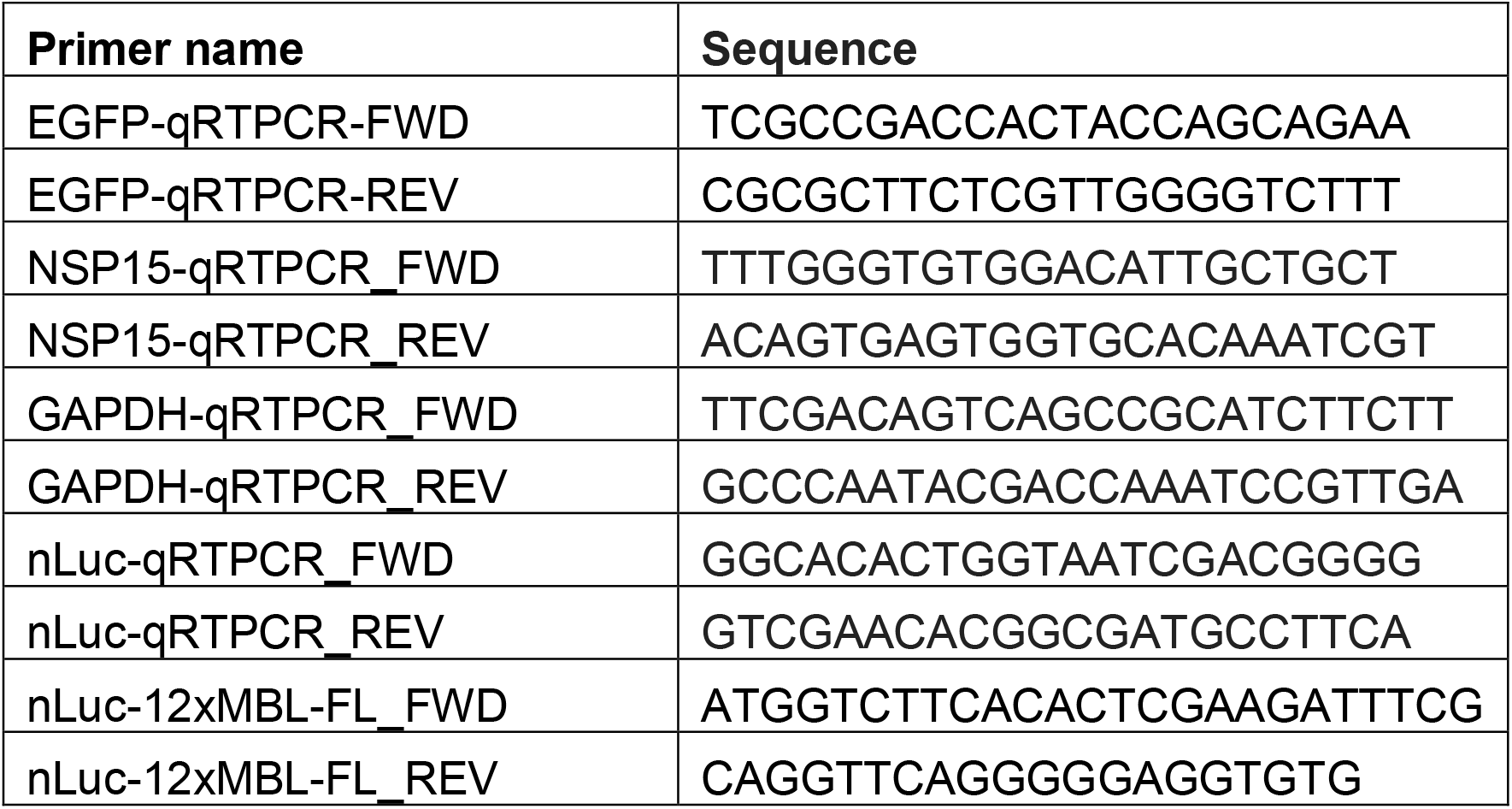

### NanoLuciferase VLP Measurement

NanoLuciferase (nLuc) was measure following manufacturors instructions. Briefly, cells were plated on a 96-well white-walled plate, infected or transfected with VLPs and after incubation the supernatant was removed, the cells were gently washed with PBS. Then, 25uL of the nLuc reagent (1:50 in manufacturor’s buffer) was added to the cells and they were incubated for 3 minutes and the signal was read on the BMG Clariostar Fluorescent plate reader. To read secreted nLuc, 20uL of the supernatant was treated with or without PK for 10 minutes at RT then 20uL of nLuc reagent was added, incubated for 3 minutes and the signal was read.

### Virus growth and purification

SARS-Related Coronavirus 2 Isolate USA-WA1/2020 was grown in VeroE6-TMPRSS2 to passage 3, titrated by plaque assay in VeroE6-TMPRSS2 with 0.75% Carboxymethyl cellulose (Sigma-C4888) overlay for 72 hrs and stained with 1% crystal violet. For experiments, the MOI was calculated and virus was added to HEK293T-Ace2-TMPRSS2 cells (a gift from Jennifer Doudna) expressing nanoLuciferase for the indicated times and treatments and the samples were again titrated by plaque assay and RNA was extracted with TRIZOL. To read nLuc in the supernatant, the supernatant was centrifuged at 10,000xg for 20 minutes and treated with 1%triton, protease K or nothing and then incubated with the nLuc reagent for 3 minutes and luminescence was read on a Promega GloMax Discover plate reader.

Dengue virus (16681) and ZIKV-PRV were both passaged on C636 cells and titrated on BHK21 cells following established protocols(38). HEK-293 cells expressing nanoLuciferase were infected with DENV2 or ZIKV at MOI=0.5. Supernatants were collected at 24 and 48 hrs and filtered through a 0.45uM PES filter. 20uL of the supernatants was added to a 96-well white walled plate and treated with PK for 10 minutes and incubated with the nLuc reagent and luminescence was measured on the BMG Clariostar Fluorescent plate reader.

### Protease Protection Assay

For the protease protection assays, 20uL of concentrated VLP or SARS-CoV2 virus supernatant or mock supernatant were incubated with or without 1% triton for 30 minutes at RT, then treated with or without 2mg/mL of protease K for 10 minutes at RT. The nLuc reagent was added for 3 minutes and luminescence read.

### siRNA treatments

The siRNAs used were synthesized by Millipore Sigma for the RdRP siRNA (35) and the Negative control #1 from ThermoFisher was used as a control. For siRNA treatment, HEK293T-Ace2-TMPRSS2 cells were transfected with 100uM of siRNAs and with Lipofectamine2000 (Thermo Fischer) along with the nanoLuciferase plasmid and incubated for 24 hrs. Then cells were infected with SARS-CoV2 MOI=0.5. The cells and supernatant were collected for nLuc reading, plaque assay and RNA extraction.

## Figure legends

**Supplemental Figure 1. Pseudo-infection of SCOV2-VLPs with luciferase-PS9**. (A) Schematic of VLP system with luciferase-PS9 adapted from (12). (B) Titration of SCOV2-VLPs with luciferase-PS9 in HEK293T-ACE2/TMPRSS2 and HEK293T cells.

**Supplemental Figure 2. Secretion of nanoLuciferase from SARS-CoV2 infected cells**. (A) A549-ACE2 cells were transfected with nLuc and infected with SARS-CoV2 for 24 hrs and the cells and supernatant were directly read for nLuc signal. *Associated with Figure 3A*. (B) HEK293 cells were transfected with nLuc and infected with DENV2 or ZIKV. The supernatant was collected at 24hrs, filtered treated with PK and nLuc signal measured. *Associated with Figure 3D*. (C-D) HEK293T-ACE2/TMPRSS2 cells were treated with siRNAs targeting the RdRp or a non-targeting control. The cells were subsequently infected with SARS-CoV2, MOI=0.5 for 48 hrs. The supernatant was collected and processed for viral genome equivalents, and infectious virus via plaque assay. The log values are plotted here. *Associated with Figure 4B*. One-way ANOVA with multiple comparisons test was performed: ns=not significant, *=p<0.05, **=p<0.01, ***p<0.001, ****p<0.0001; n=3.

## Author Contributions

RCG and JF conceptualized the project, RCG carried out the experiments, RCG and JF wrote the manuscript.

