## Supplementary figures and images for "SARS-CoV2 associated secretion of nanoLuciferase reports on virus and Virus-Like Particle production"

### Supplemental Figures

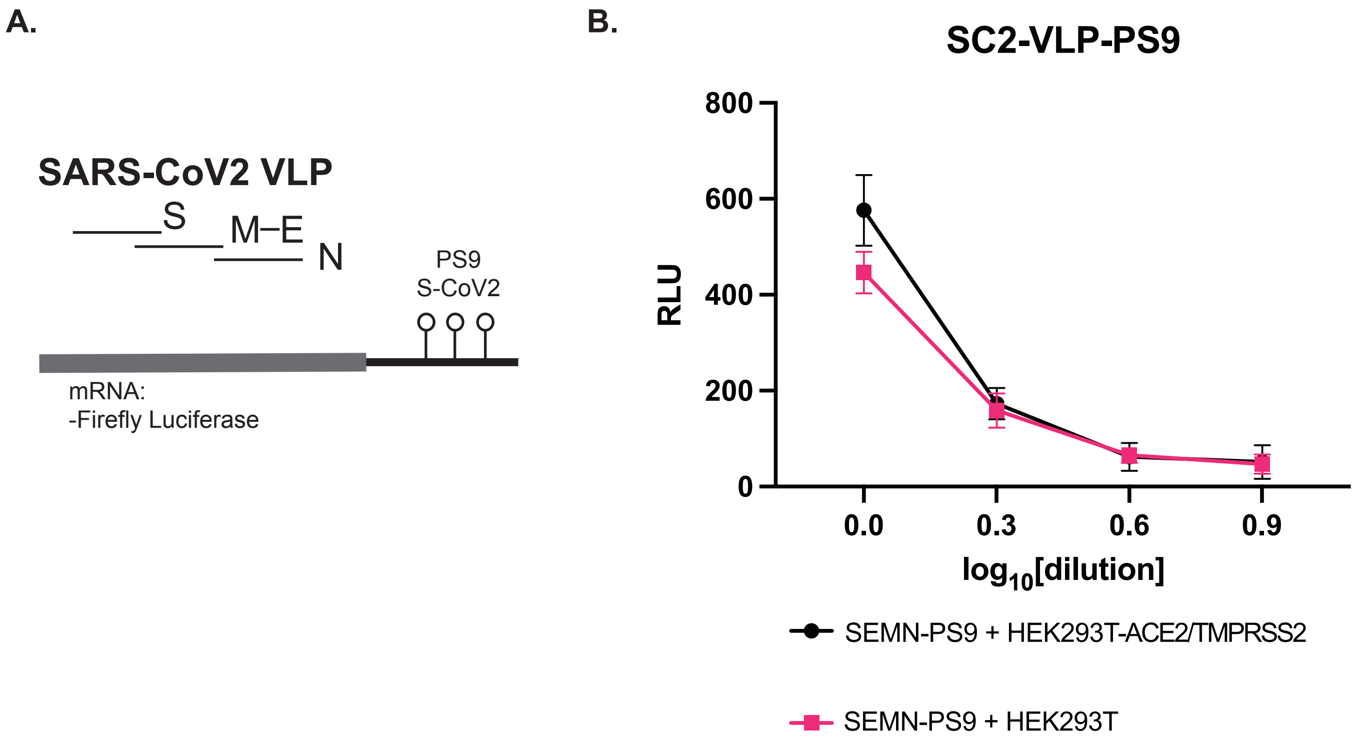

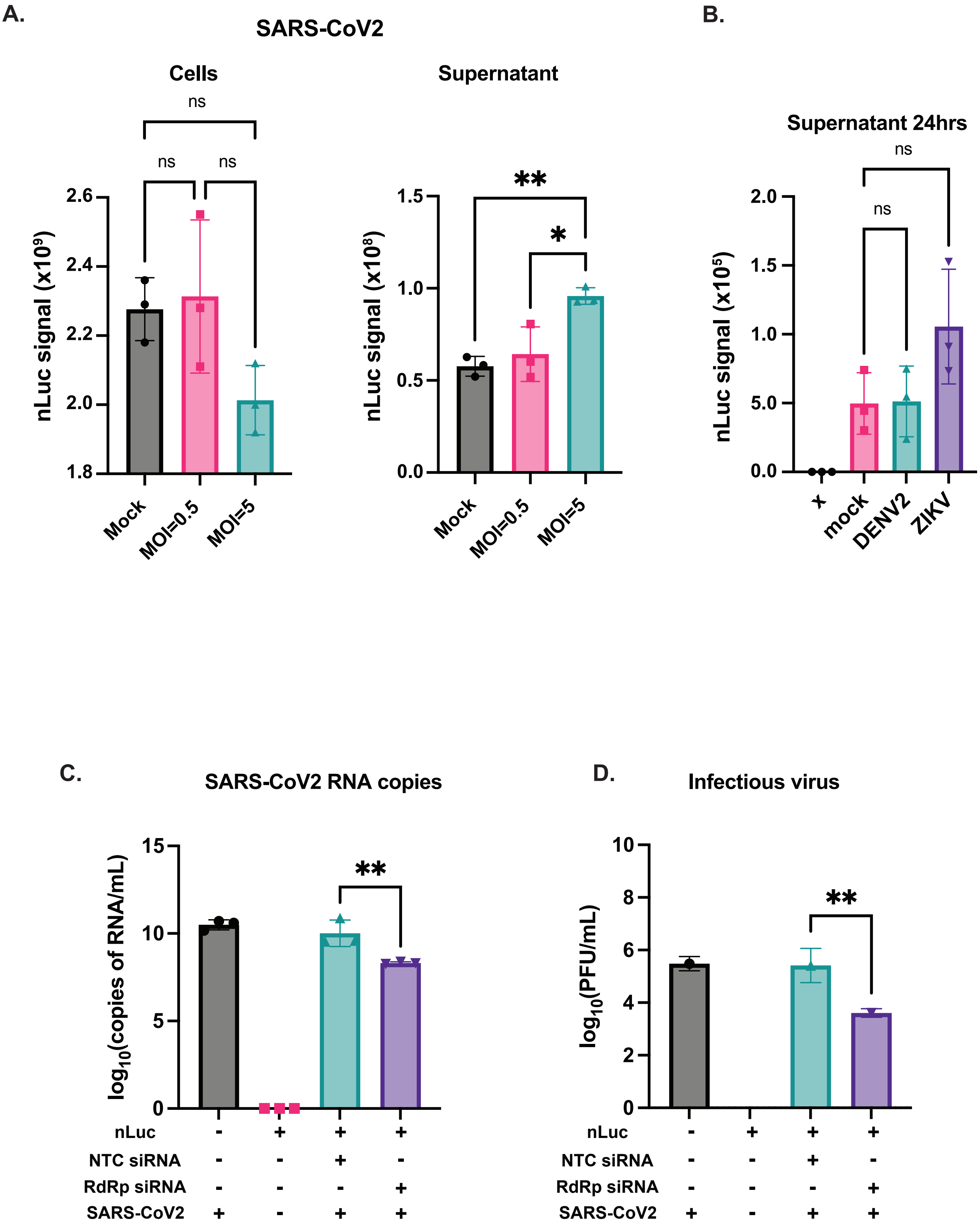
